# Survival on a semi-arid island: submersion and desiccation tolerances of fiddler crabs from the Galapagos Archipelago

**DOI:** 10.1101/2020.05.27.120014

**Authors:** Mariana V. Capparelli, Carl L. Thurman, Paloma Gusso Choueri, Denis Moledo Abessa, Mayana Karoline Fontes, Caio Rodrigues Nobre, John Campbell McNamara

## Abstract

During tidal cycles, semi-terrestrial fiddler crabs are subject to alternating periods of submersion and desiccation. Here, we compare physiological and biochemical adjustments to forced submersion and desiccation in two fiddler crabs from the Galapagos archipelago: the indigenous *Leptuca helleri*, and *Minuca galapagensis*. We examine ecological distributions and habitat characteristics using transect analysis; survival after 6 h forced submersion at different salinities (0, 21 and 42 ‰S), and after 6 or 12 h desiccation challenge, including alterations in hemolymph osmolality; and, oxidative stress responses in the gills and hepatopancreas, accompanying glutathione peroxidase (GPx), glutathione S-transferase (GST) and glutathione reductase (GR) activities, and lipid peroxidase (LPO). We provide an integrated biomarker response (IBR) index for each species based on oxidative stress in each tissue and condition. Our transect study revealed that *L. helleri* occupies an intertidal niche while *M. galapagensis* is supralittoral, *L. helleri* being less resistant to submersion and desiccation. After 6 h submersion, *L. helleri* survived only at 21 ‰S while *M. galapagensis* survived at all salinities. Hemolymph osmolality decreased at 0 ‰S in *M. galapagensis*. After 6 h desiccation, osmolality decreased markedly in *L. helleri* but increased in *M. galapagensis*. Enzyme assays were not performed in *L. helleri* owing to high mortality on submersion/desiccation challenge. After submersion in *M. galapagensis*, hepatopancreas GPx activities decreased in 0 and 21 ‰S while GR activity was strongly inhibited at all salinities. Gill LPO decreased in 42 ‰S. On desiccation in *L. helleri*, GPx activity was inhibited in the hepatopancreas but increased in the gills. GST activity increased while LPO decreased in both tissues. After desiccation in *M. galapagensis*, hepatopancreas GPx activity increased. Both hepatopancreas and gill GST and GR activities and LPO were strongly inhibited. The IBR indexes for *L. helleri* were highest in fresh caught crabs, driven by gill and hepatopancreas LPO. For *M. galapagensis*, submersion at 21 ‰S contributed most to IBR, LPO in both tissues responding markedly. *Leptuca helleri* appears to be a habitat specialist adapted to a narrow set of niche dimensions while *M. galapagensis* survives over a much wider range, exhibiting little oxidative stress. The species’ physiological flexibilities and limitations provide insights into how fiddler crabs might respond to global environmental change on semi-arid islands.

## Introduction

Fiddler crabs are typical inhabitants of the estuarine environments of tropical and temperate zones (Crane, 1975). Many biological factors such as vegetation cover and resource competition, and abiotic attributes like edaphic characteristics, sediment grain size, organic content, and salinity and temperature influence their distribution (Nobbs, 2003; Thurman et al., 2013; Mokhtari et al., 2015; Checon and Costa, 2017). Such heterogeneous habitats range from salt marshes and exposed, dry sandy beaches to shaded, muddy mangrove forests (Thurman, 1984; Thurman et al., 2013), which can be physiologically challenging to fiddler crabs (Allen and Levinton, 2014; Munguia et al., 2017, Thurman et al., 2017). During tidal cycles, many fiddler crab species are exposed to temperature and salinity variation (Helmuth et al., 2006; Schneider, 2008; Somero, 2002) and face desiccation (Allen et al., 2012; Chapman and Underwood, 1996; Miller et al., 2009; Thurman, 1998).

Nevertheless, fiddler crabs are resilient components even of degraded ecosystems and some can tolerate severely polluted habitats (Capparelli et al., 2016). Certain species are more generalist in their ecological demands, inhabiting a diversity of environments while others are more specialized, exhibiting restricted habitat preferences. Given predicted alterations to mangrove communities, owing to habitat loss and climate change (Saintilanet al. 2014), some fiddler crab species may not be able to survive in novel habitats, despite their tolerances of variation in ambient parameters. Many minimize the effects of desiccation, for example, by inhabiting burrows as a refuge (Powers and Cole, 1976), albeit with fewer opportunities to forage and reproduce (Allen and Levinton, 2014). Desiccation and submersion tolerance varies among fiddler crabs and is an indicative of their resistance to aerial exposure during low tide (Thurman, 1998; Levinton et al., 2015; Principe et al., 2018). Large scale landscape changes, such as mangrove deforestation or altered ocean levels, may cause physiological changes resulting from submersion and desiccation challenge that affect ecological distribution patterns (Araújo et al., 2007, Wilson et al., 2005) and survival.

Fiddler crab ecology has been of much interest (Landstorfer et al., 2010, Cuellar-Gempeler and Munguia, 2013; Costa and Soares-Gomes, 2015, Thurman et al. 2003, 2005, 2017), but the effects of desiccation and submersion on their physiology is poorly known (Levington et al., 2015). Species distributed within the upper tidal zone like *Minuca rapax* are more resistant to dissection than intertidal species such as *Leptuca thayeri* (Principe et al., 2018). Osmoregulatory ability and aerial exposure show no general trend (Gilles and Péqueux, 1983; Borecka et al., 2016) although aerial exposure can increase hemolymph osmolality due to dehydration, physiological responses varying among species (Thurman, 1998).

Aerial exposure represents a major challenge for intertidal organisms, since it leads to dehydration and metabolic stress (Allen et al., 2012; Chapman and Underwood, 1996; Miller et al., 2009). Cellular oxidative stress occurs when the rate of production of reactive oxygen species (ROS) exceeds their decomposition by antioxidant systems, increasing oxidative damage such as lipid peroxidation, enzyme inactivation, DNA base oxidation and protein degradation (Halliwell, 1993; Lemaire and Livingstone, 1993). Cellular protection against ROS includes the glutathione system of specific antioxidant enzymes such as glutathione peroxidase (Sies et al., 1979; Keeling and Smith, 1982; Sies, 1993) and glutathione S-transferases (Tan et al., 1987), together with complementary enzymes like glutathione reductase that produce glutathione and NADPH, maintaining cellular antioxidant status (Reed, 1986). Antioxidant enzymes and oxidative damage levels as indicators of oxidative stress have been evaluated in *M. rapax* from contaminated environments (Capparelli et al., 2019). However, little is known with regard to modulation of oxidative stress in fiddler crabs from pristine habitats during submersion and desiccation challenge.

The two fiddler crab species found on the Galapagos Archipelago, *M. galapagensis* Rathbun 1902 and *L. helleri* Rathbun 1902, were originally considered endemic (Rathbun, 1902), and their ecological preferences are very sketchy. Boone (1927) quoting Beebe (1924) stated that *M. galapagensis* typically occupied “salt marshes and tidal flats”, with burrows at the high-tide mark of about 2 cm diameter and 20-30 cm deep. Garth (1948) described *M. galapagensis* as burrowing near brackish-water lagoons or on mud flats with iron-red or gray colored substrate; *L. helleri* was usually found on sandy-mud among mangrove roots, the species supposedly separated by their habitat preferences. Von Hagen (1968) considered *M. galapagensis* as very flexible in occupying habitats with widely differing substrates. Peck (1994) reported *M. galapagensis* to inhabit soft mud in mangroves, and *L. helleri* to occur on sandy or muddy intertidal flats, with no ecological overlap.

The present study aims to compare the effect of submersion and desiccation challenge in *L. helleri* and *M. galapagensis* from mangrove areas on Santa Cruz Island, in the Galapagos Archipeligo. We hypothesize that *L. helleri*, an ecologically demanding species, would be more sensitive to submersion and desiccation than *M. galapagensis*, a generalist species, such tolerances subsidizing their ecological preferences. Understanding the differential effects of extreme conditions of desiccation and submersion on fiddler crabs from areas with different degrees of exposure to water helps to predict how these species behave in possible environmental change scenarios. Specifically we evaluate the species’: (1) density distributions as a function of selected ambient parameters; (2) tolerance of submersion and desiccation, and hemolymph osmoregulatory ability; and (3) oxidative stress response to submersion and desiccation.

## Materials and methods

### Study area

The study area was located in a stretch of arid lowland on the southern coast of Santa Cruz Island, one of the 13 major islands that constitute the Galapagos Archipelago, Ecuador. Santa Cruz Island is the second largest in the archipelago, is 986 km^2^ in area and has a maximum altitude of 864 m. Recent temperatures and precipitation have ranged from lows of 21-22 °C (mean 22 °C) and 0.5 mm in September 2018 to highs of 25-29 °C (mean, 27 °C) and 64.9 mm in April 2019 (www.worldweatheronline.com/santa-cruz-weather-averages/galapagos/ec.aspx). Around 90% of the terrestrial area is protected as part of the Galapagos National Park (Servicio Parque Nacional Galapagos, 2006; Moity et al., 2019).

### Study organisms and ecological transects

The two fiddler crab species found on the Galapagos Archipelago are *Minuca galapagensis*, and the endemic *Leptuca helleri* (Rathbun, 1902). Originally placed in the genus *Uca*, they now belong to separate genera: *Minuca* (Bott) and *Leptuca* (Bott) (Shih et al., 2016). There is no information on the conservation status of either species in the IUCN Red List of Threatened Species (https://www.iucnredlist.org, May, 2020).

Potential habitats on Santa Cruz Island were reconnoitered for fiddler crab colonies and areas for transect analyses were selected for *Leptuca helleri and Minuca galapagensis* based on salinity, abundance and ease of access. Most sites had only one species.

Among the eight possible locations, the best site for characterizing *L. helleri* was on the eastern edge of the Playa de los Alemanes (00.75243° S, 90.31087° W), a beach consisting of coarse coralline sand with sea shells and broken coral, near a mangrove stand *(Rhizophora* sp.) (Figures 1 and 2). For *M. galapagensis*, the optimum area was a large sand flat named “Tortuga Flats” by us, enclosed by mangroves on the northern shore of Bahia Tortuga, a fine white sand beach (00.76174° S, 90.34111° W). The transects were established orthogonally to the nearest water source. The “Tortuga Flats” transect ran 30 m inland from the supralittoral sand berm towards a large shallow pond while the Playa de los Alemanes transect ran 16 m down the beach from the supralittoral zone to the low water mark (Figures 1 and 2).

**Figure 1.**
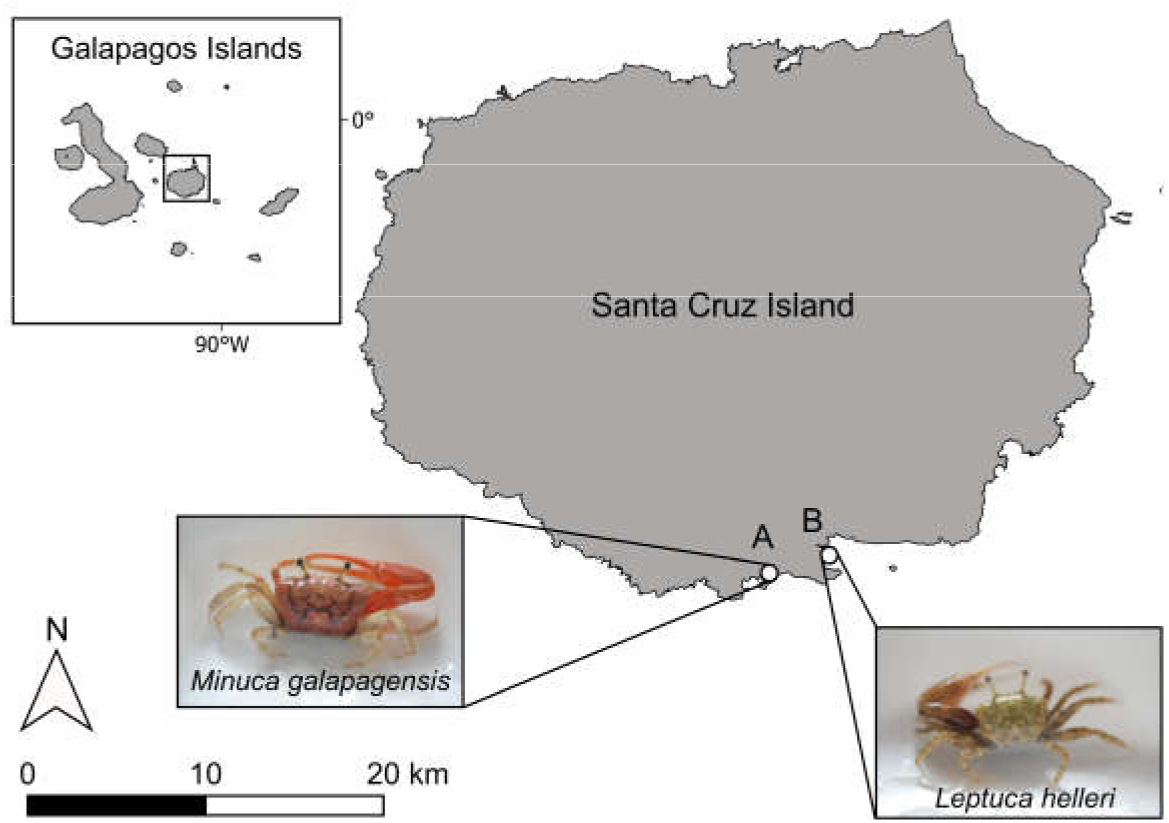
The study area. Santa Cruz Island is one of the 13 main islands that form the Galapagos Archipelago, Ecuador. It is the second largest island and has an area of 986 km^2^ and a maximum altitude of 864 m. Approximately 90% of the island is protected as part of the Galapagos National Park. The Galapagos fiddler crabs *Leptuca helleri* and *Minuca galapagensis* abound in the coastal mangrove ecosystems. A, Bahia Tortuga collecting site (00.76174° S, 90.34111° W). B, Playa de los Alemanes collecting site (00.75243° S, 90.31087° W).

**Figure 2.**
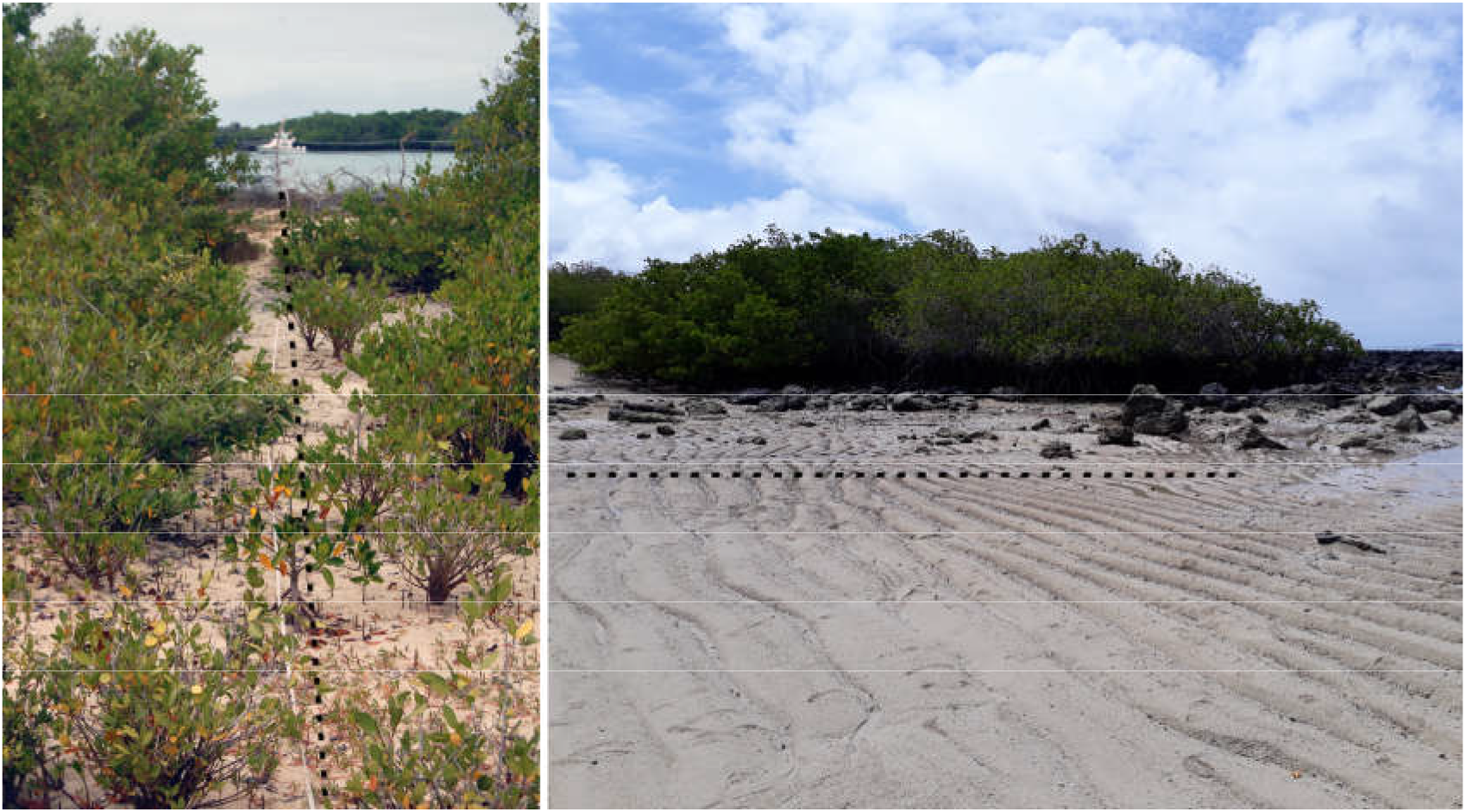
Sites of transect analyses for the Galapagos fiddler crabs. For *Minuca galapagensis* (left panel), the optimum area was an extensive, sheltered sand flat (“Tortuga Flats”) amongst the mangroves behind the sand berm on the shore of Bahia Tortuga (00.76174° S, 90.34111° W). For the endemic species *Leptuca helleri* (right panel), the best area was adjacent to an exposed mangrove stand on the eastern edge of Playa de los Alemanes (00.75243° S, 90.31087° W). Dotted lines indicate approximate transects taken orthogonally to the nearest water source.

Vegetation coverage and the number of crab burrows in 1 m^2^ quadrats were recorded at 2 m intervals along the transects. On the *M. galapagensis* transect, percentage soil moisture was taken at a depth of 15 cm with an electronic soil probe. Soil temperatures were measured on the substrate surface and at 10 cm depth. Each burrow was excavated, the inhabitant identified, a water sample collected, and the water table depth measured. The osmolality of the water samples (in mOsm/kg H_2_O) was measured using a vapor pressure micro-osmometer (Wescor 5520, Logan, UT).

### Crab collections

Adult, intermolt specimens of *M. galapagensis* and *L. helleri* of either sex were collected in June and July 2019 (dry season) from the “Tortuga Flats” at Bahia Tortuga (00.76174° S, 90.34111° W) and the sandy beach at Playa de los Alemanes (00.75243° S, 90.31087° W), Santa Cruz Island, Galapagos (see Figures 1 and 2) (collecting permit #083-2019 from the Dirección del Parque Nacional Galápagos to MVC). Only *L. helleri* occurred at Playa de los Alemanes while both species were abundant at “Tortuga Flats”.

The crabs were transported to the Fabricio Valverde laboratory at the Charles Darwin Research Station in plastic boxes containing sponge cubes moistened with seawater from the collecting sites. Only non-ovigerous, intermolt crabs of carapace width greater than 10 mm for *M. galapagensis* and 3 mm for *L. helleri* were used.

To acclimatize to laboratory conditions before use, the crabs were maintained unfed for 2 days after collection at 25 °C under a 12 h light: 12 h dark natural photoperiod, with free access to a dry surface, in plastic boxes containing seawater from the respective collection sites (29 ‰S).

### Submersion and desiccation protocols

To examine the effects of forced submersion, crabs were maintained fully submerged, simulating the high tide covering their burrows. Groups of 10 crabs each were submerged for 6 h at salinities of 0 ‰S [distilled H_2_O, hypo-osmotic medium], 21 ‰S [isosmotic reference medium] or 42 ‰S [hyper-osmotic medium] in individual plastic jars containing 250 mL of medium. Saline media were prepared by diluting seawater with bottled water or adding Instant Ocean sea salts. Salinities were checked using a hand-held refractometer (American Optical Company, MA). Six hours is roughly the period a crab would be submerged naturally between the pre- and post-high tide (Batista, 2010; Capparelli et al., 20017).

To examine the effects of desiccation, previously blotted crabs were held in individual dry containers for 6 or 12 h. Mortality was totaled at the end of both experiments. Crabs were considered dead if they could not right themselves.

After the submersion and desiccation experiments, the crabs were cryo-anesthetized in crushed ice for 10 min. A hemolymph sample was then drawn through the arthrodial membrane at the base of the posterior-most pereiopod into a 1 mL syringe, and all gills and the hepatopancreas were dissected, placed in individual, labeled micro-Eppendorf tubes and frozen at −80 °C for posterior analysis. Two-day acclimatized crabs, dissected at the beginning of the experiments (Time = 0 h), were used as reference control crabs (fresh caught crabs).

Samples were transported in dry ice by air to laboratories in Brazil. The entrance of samples into Brazil at Guarulhos Airport was authorized by the Ministério da Agricultura, Pecuária e Abastecimento (licence #000014.0020214/2019 to JCM).

### Measurement of hemolymph osmolality

Hemolymph osmolality in both crab species was measured in 10 μL aliquots or occasionally employing 2-3 hemolymph pools, using a vapor pressure micro-osmometer (Wescor 5500, Logan, UT).

### Oxidative stress assays

Oxidative stress activities were measured in hepatopancreas and gill homogenates from *M. galapagensis* and *L. helleri*. Immediately before the assays, samples were thawed on ice and homogenized in a Tris-HCl buffer solution (4% w/v, in mmol L^-1^, Tris 50, Ethylenediamine tetra acetic acid 1, Dithiothreitol 1, Sucrose 50, KCl 150, Phenylmethylsulfonyl fluoride 1, pH 7.6). Aliquots were separated to analyze lipid peroxidation (LPO). The remaining homogenates were centrifuged (Eppendorf model 5804R, Eppendorf North America, Hauppauge, NY) at 12,000 × g and 4 °C for 20 min, and the supernatants used for the enzyme assays.

Glutathione S-transferase (GST) and glutathione peroxidase (GPx) and glutathione reductase (GR) activities were assayed following the protocols described by Keen et al. (1976), Sies et al. (1979) and Mcfarland et al. (1999), respectively. All enzyme activities were measured spectrophotometrically at 340 nm. LPO analyses were performed by fluorescence spectroscopy (excitation 516 nm, emission 600 nm) using the thiobarbituric acid reactive substances (TBARS) method (Wills, 1987).

All assays were performed using a BioTek Synergy HT Multi-Detection Microplate Reader (BioTek, Winooski, VT). Enzyme activities were normalized by total protein content, measured using Bradford’s (1976) method.

### Integrated biomarker response indexes

The overall effects of 6-h forced submersion at the different salinities or desiccation for 6 or 12 h on biomarker activities in each tissue were quantified using integrated biomarker response indexes (Beliaeff and Burgeot, 2002; Liu et al. 2013; Perussolo et al., 2019). To create each index, each individual biomarker response value was normalized by subtraction from the grand mean value for all replicates and divided by its standard deviation. Each normalized value was then added to the minimum absolute value obtained for each biomarker (Z score). Each mean biomarker Z score was then multiplied by a weighting (GST, GPx, GR = 1; LPO = 2), designated according to the systemic importance each activity (biomarker response score). To obtain the integrated biomarker response index (IBR), the sum of all biomarker response scores for each condition and tissue was then divided by the sum of all weightings, providing the final degree of effect for each condition.

Biomarker response scores were then plotted as Radar charts using Microsoft Excel software (Microsoft Corp., Redmond, WA, USA).

### Statistical Analyses

After verifying the data for normality of distribution and homoscedasticity by applying the Shapiro-Wilks and Bartlett tests, respectively, the effect of salinity or of desiccation time on hemolymph osmolality and enzyme activities was evaluated using one-way analyses of variance. Differences between means within each parameter were located using the Student-Newman-Keuls *post-hoc* multiple comparisons procedure. A minimum significance level of P = 0.05 was employed throughout.

## Results

### Ecological transect sampling

Figure 3 shows the variation in burrow densities and in abiotic parameters like interstitial moisture content, salinity and burrow temperature measured along the respective transects taken at the Playa de los Alemanes and Bahia Tortuga collecting sites where the fiddler crabs *L. helleri* and *M. galapagensis* were abundant. The two species were sympatric at the Bahia Tortuga site.

**Figure 3.**
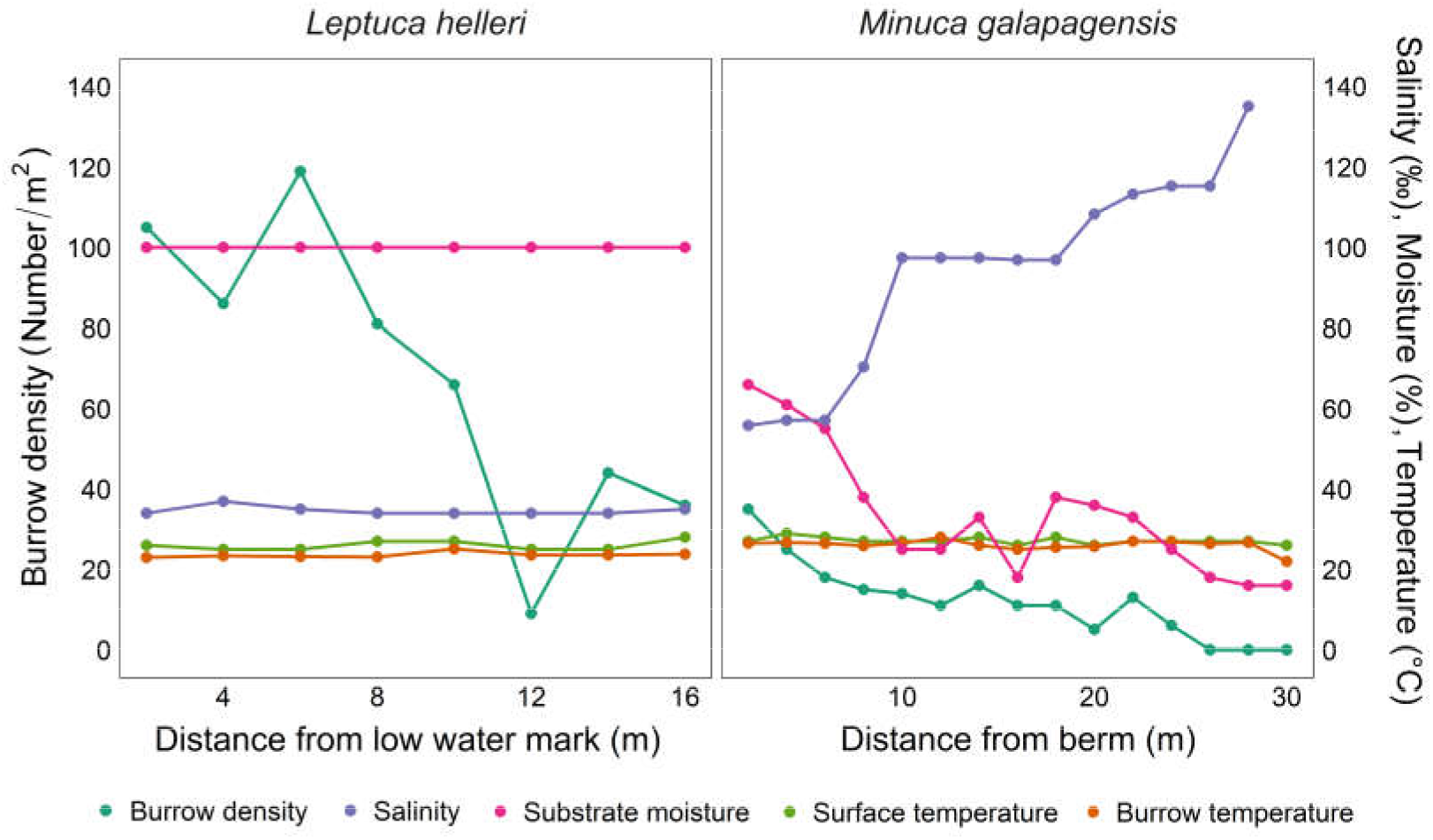
Variation in burrow densities and abiotic parameters (salinity ‰S, substrate moisture %, and surface and burrow temperatures °C) measured along the respective transects made at the Playa de los Alemanes (00.75243° S, 90.31087° W) (fine white sand) and Bahia Tortuga (00.76174° S, 90.34111° W) (coarse coralline sand with sea shells and broken coral) sites on Santa Cruz Island, Ecuador, where the fiddler crabs *Leptuca helleri* and *Minuca galapagensis* were abundant.

*Leptuca helleri* showed the highest burrow densities (120 burrows/m^2^) closest to the low water mark. For *M. galapagensis*, burrow density was highest near the shoreline berm (40 burrows/m2). Burrow densities decreased progressively along the transects to minima of 36 and 6, respectively, at 16 and 24 meters from the low water mark and berm. Burrow salinity (34 to 37 ‰S), temperature (23.0 to 25.1 °C) and moisture (100%) were fairly constant for *L. helleri*. However, for *M. galapagensis*, burrow moisture decreased from 66 to 25% while salinity increased from 56 to 115 ‰S along the transect; burrow temperature ranged from 25.0 to 28.0 °C. Substrate surface temperatures were 26 to ≈29 °C for both species while burrow temperatures (23 to 25 °C) were slightly less for *L. helleri*.

### Effects of forced submersion and desiccation

#### Survival

*Minuca galapagensis* showed no mortality in either condition (Table 2). In contrast, *L. helleri* did not survive at all on submersion at 0 and 42 ‰S. At 21 ‰S, survival was 40%. During desiccation, *L. helleri* showed 30% and 0% survival after 6 and 12 h, respectively (Table 2).

**Table 2.**
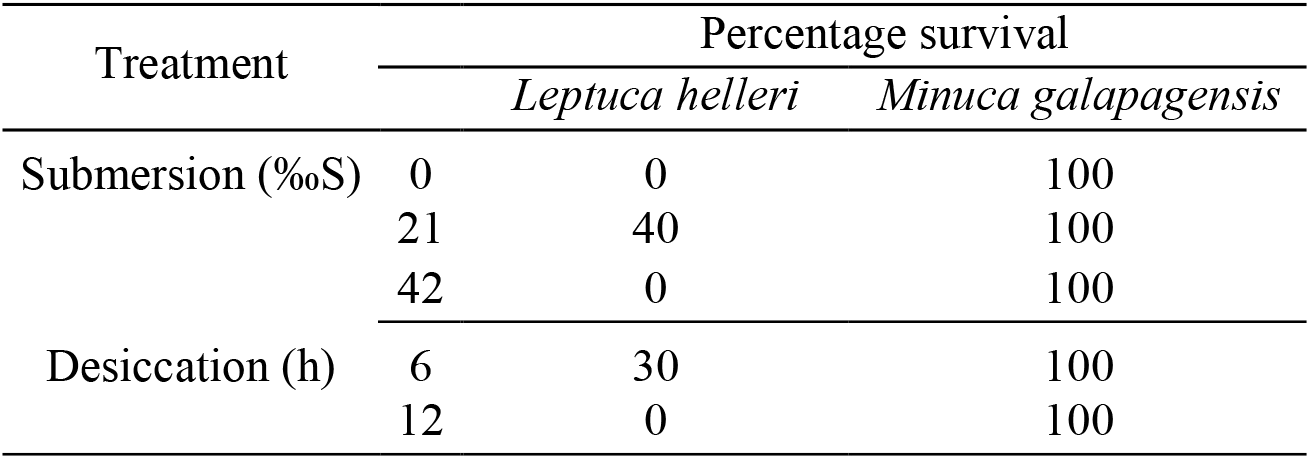
Percentage survival of the Galapagos fiddler crabs *Leptuca helleri* and *Minuca galapagensis* from Santa Cruz Island, Ecuador, during forced submersion for 6 h at different salinities and up to 12 h desiccation.

#### Hemolymph osmoregulatory ability

Hemolymph osmolality in *L. helleri* was unchanged after 6 h of forced submersion at 21 ‰S compared to fresh caught control crabs (Figure 4) and was strongly hyper-regulated (Δ = +405 mOsm/kg H_2_O). In *M. galapagensis*, osmolality decreased after 6 h at 0 ‰S compared to fresh caught controls while at 21 and 40 ‰S osmolality was unaltered. In these salinity-challenged crabs, hemolymph osmolality increased progressively and was hyper-regulated (Δ = +230 mOsm/kg H_2_O) in 21 ‰S but hypo-regulated (Δ = −310 mOsm/kg H_2_O) in 42 ‰S (Figure 4).

**Figure 4.**
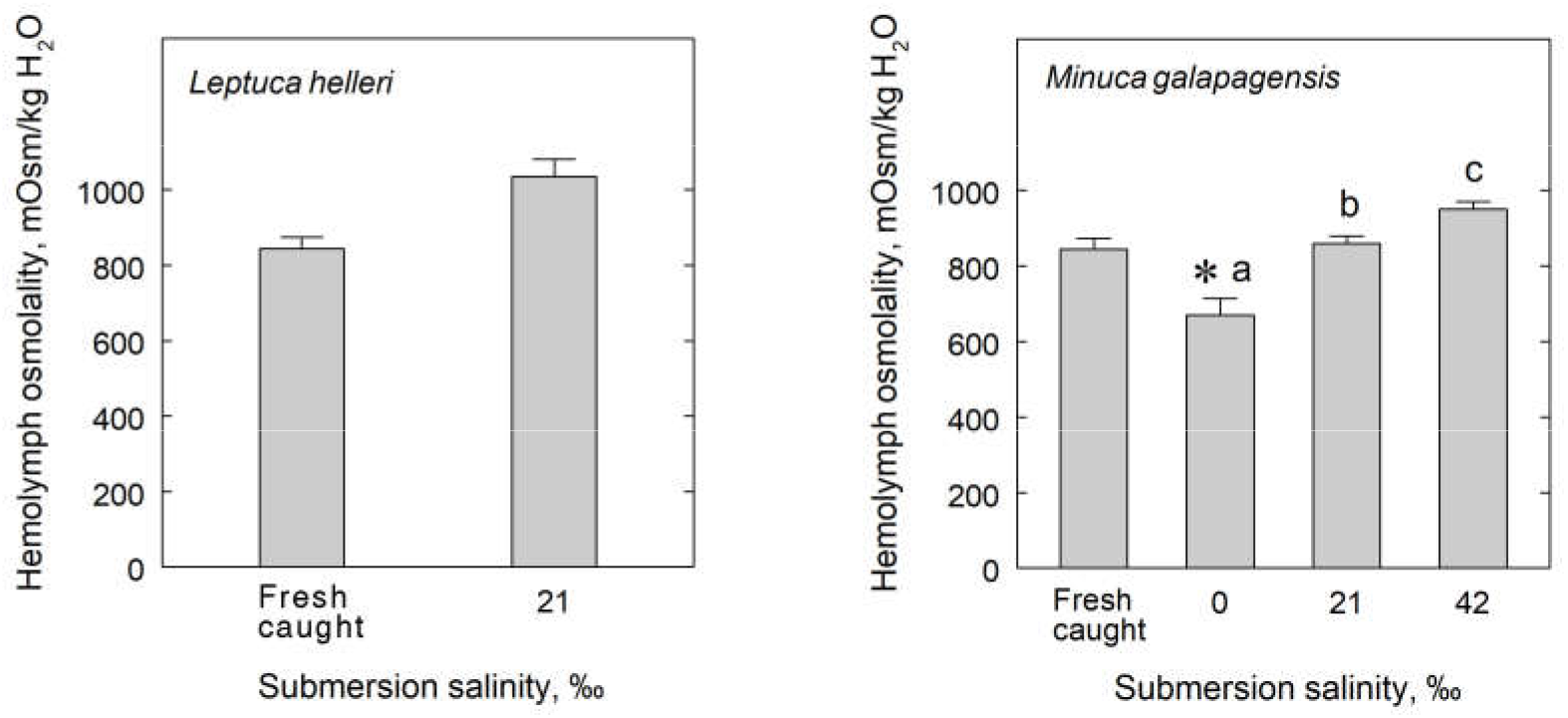
Osmoregulatory ability in the Galapagos fiddler crabs *Leptuca helleri* and *Minuca galapagensis* when maintained fully submerged for 6 h at different salinities (0 ‰S [distilled H_2_O, hypo-osmotic challenge], 21 ‰S [630 mOsm/kg H_2_O, isosmotic reference medium] or 42 ‰S [1,260 mOsm/kg H_2_O, hyper-osmotic challenge) after 2 days held at 29 ‰S (670 mOsm/kg H_2_O, fresh caught crabs). Data are the mean ± SEM (N=10). *P<0.05 compared to fresh caught crabs; different letters indicate significantly different groups

Hemolymph osmolality in *L. helleri* decreased 0.4-fold after 6 h desiccation compared to fresh caught control crabs (Figure 5). In *M. galapagensis*, hemolymph osmolality increased 1.7-fold after 6 h, decreasing after 12 h to fresh caught control values (Figure 5).

**Figure 5.**
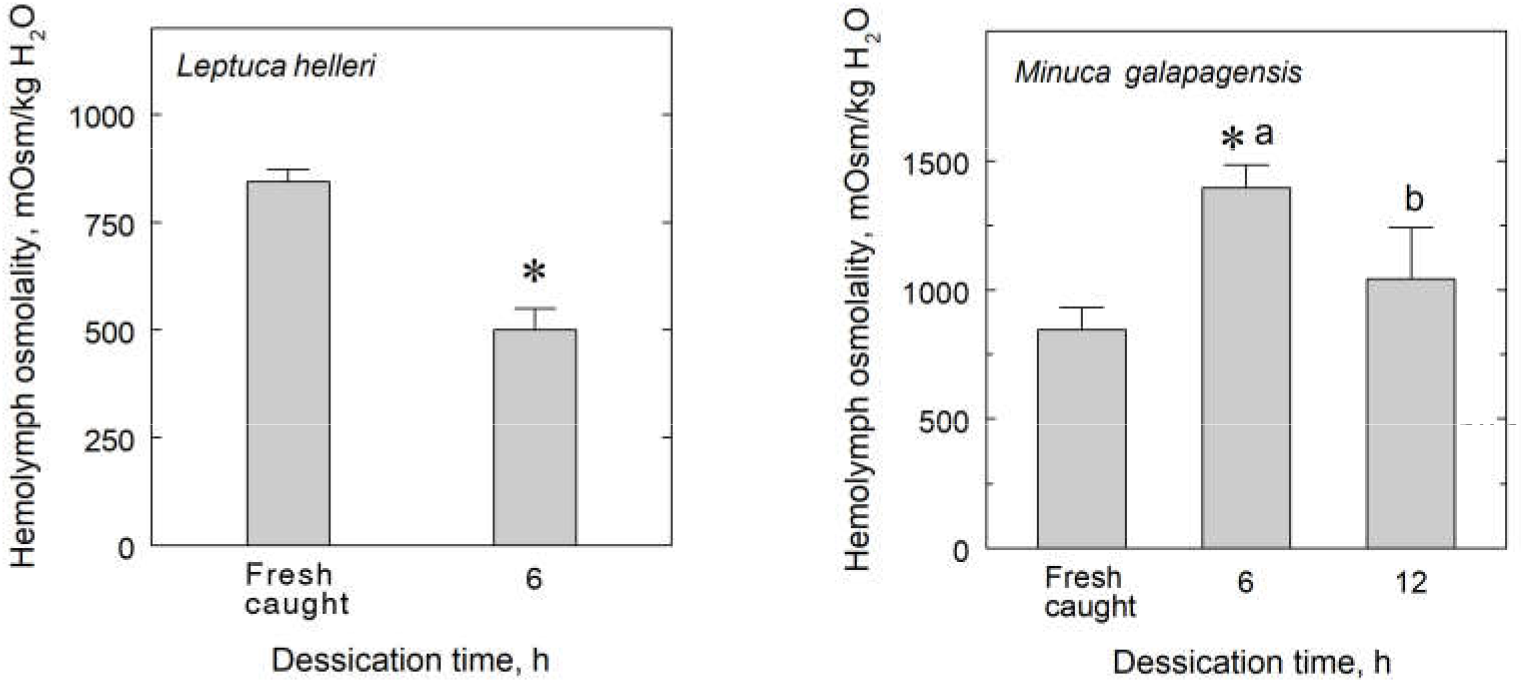
Changes in the hemolymph osmolality of the Galapagos fiddler crabs *Leptuca helleri* and *Minuca galapagensis* when kept emerged without access to water for up to 12 h, after 2 days held at 29 ‰S (670 mOsm/kg H_2_O, fresh caught crabs). Data are the mean ± SEM (N=10). *P<0.05 compared to fresh caught crabs; different letters indicate significantly different groups.

#### Oxidative stress enzymes

After forced submersion of *M. galapagensis* for 6 h, hepatopancreas GPx activities decreased in 0 and 21 ‰S compared to fresh caught control crabs (Figure 6). In 42 ‰S, activity increased to control crab levels. GST activities were unaffected compared to fresh caught crabs, although activity decreased in 42 ‰S compared to 0 and 21‰S. Hepatopancreas GR activity was strongly inhibited at all salinities (Figure 6) while LPO activities were unaffected by salinity compared to fresh caught crabs (Figure 6). However, LPO activities increased in 21 and 42 ‰S compared to 0 ‰S.

**Figure 6.**
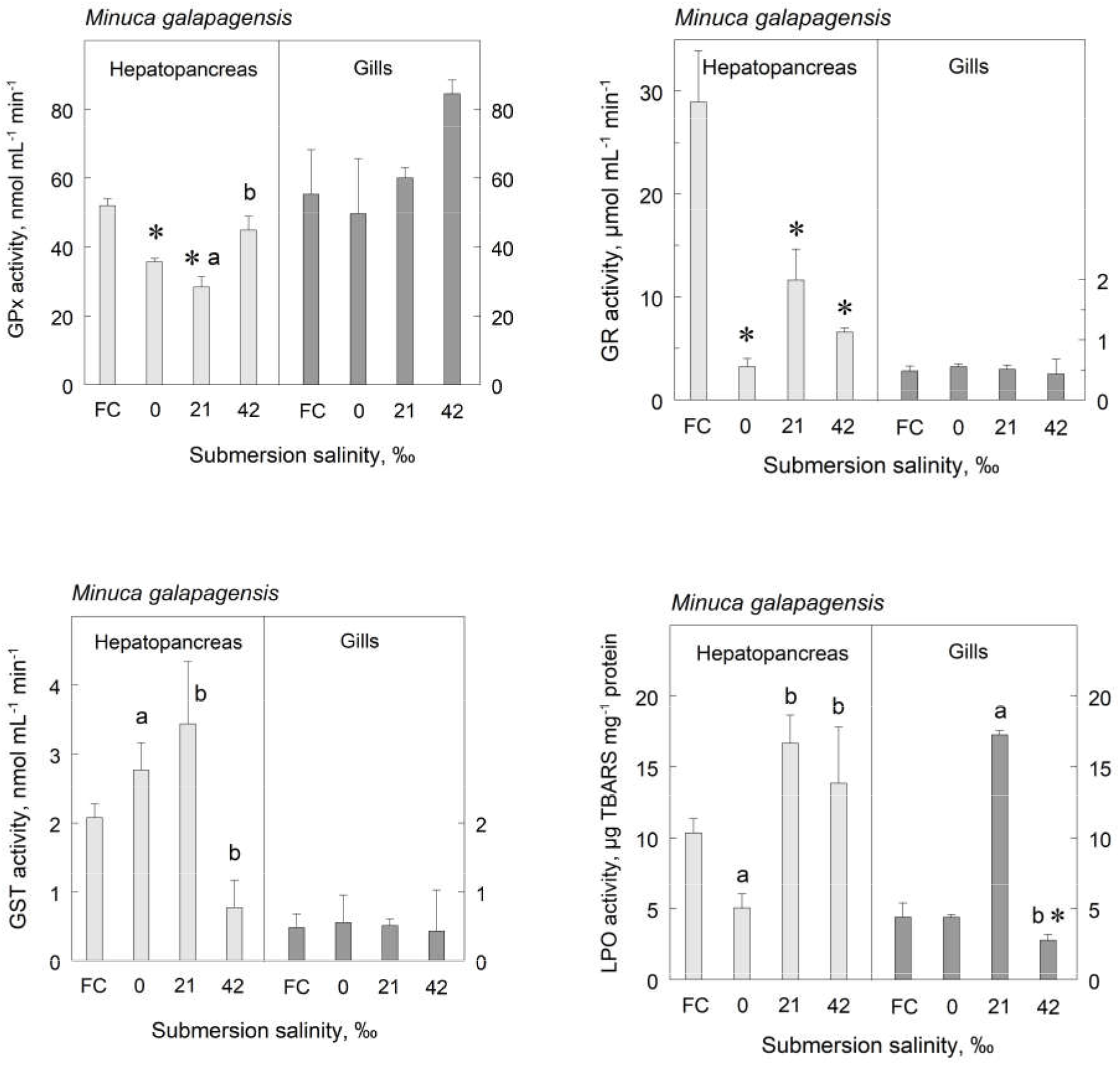
Enzyme activities (Glutathione peroxidase [GPx], Glutathione S-transferase [GST], Glutathione Reductase [GR]) and Lipid peroxidation [LPO] in hepatopancreas and gill homogenates from the Galapagos fiddler crab *Minuca galapagensis* maintained fully submerged for 6 h at different salinities (0 ‰S [distilled H_2_O, hypo-osmotic challenge], 21 ‰S [isosmotic reference medium] or 42 ‰S [hyper-osmotic challenge) after 2 days held at 29 ‰S (fresh caught crabs, FC). Data are the mean ± SEM (N=10). *P<0.05 compared to fresh caught crabs; different letters indicate significantly different groups.

There was no effect of submersion salinity on gill GPx, GST or GR activities compared to fresh caught crabs (Figure 6). Gill LPO activities increased in 21 ‰S but decreased in 42 ‰S to values below the fresh caught controls (Figure 6).

During desiccation in *M. galapagensis*, hepatopancreas GPx activity increased after 6 and 12 h compared to fresh caught control crabs (Figure 7). Gill GPx activities were unaltered (Figure 7). Both hepatopancreas and gill GST, GR and LPO activities were strongly inhibited after 6 and 12 h desiccation (Figure 7).

**Figure 7.**
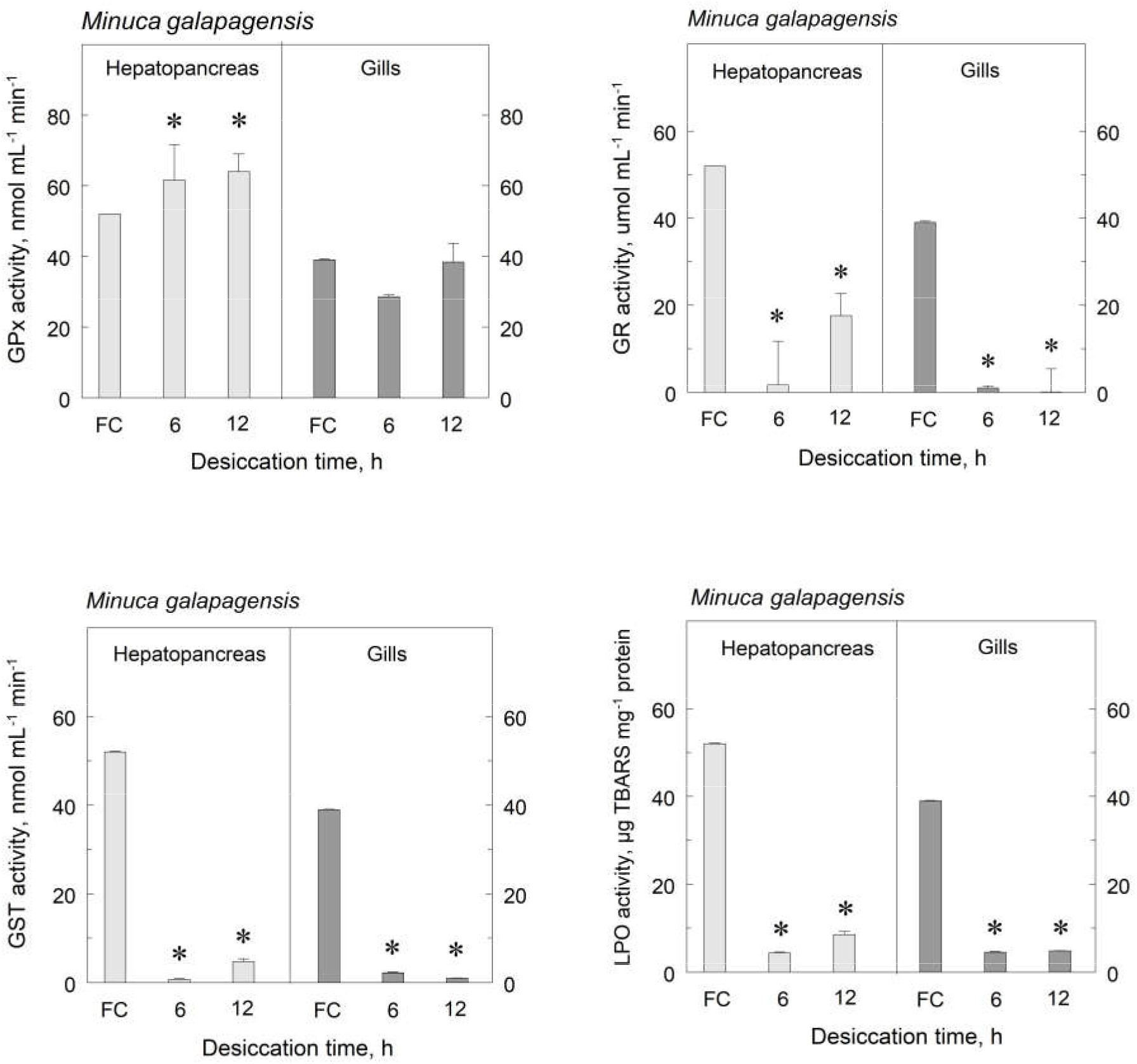
Enzyme activities (Glutathione peroxidase [GPx], Glutathione S-transferase [GST], Glutathione Reductase [GR] and Lipid peroxidation [LPO]) in hepatopancreas and gill homogenates from the Galapagos fiddler crab *Minuca galapagensis* kept emerged without access to water for up to 12 h, after 2 days held at 29 ‰S (fresh caught crabs, FC). Data are the mean ± SEM (N=10). *P<0.05 compared to fresh caught crabs.

In *L. helleri*, forced submersion resulted in high mortality (40%) and enzymatic assays could not be performed.

During desiccation, *L. helleri* survived only for 6 h. GPx activity was inhibited in the hepatopancreas but increased in the gills compared to fresh caught control crabs (Figure 8). GST activity increased in both tissues (Figure 8). GR activities were unaltered by desiccation (Figure 8) while LPO activities decreased in both tissues (Figure 8).

**Figure 8.**
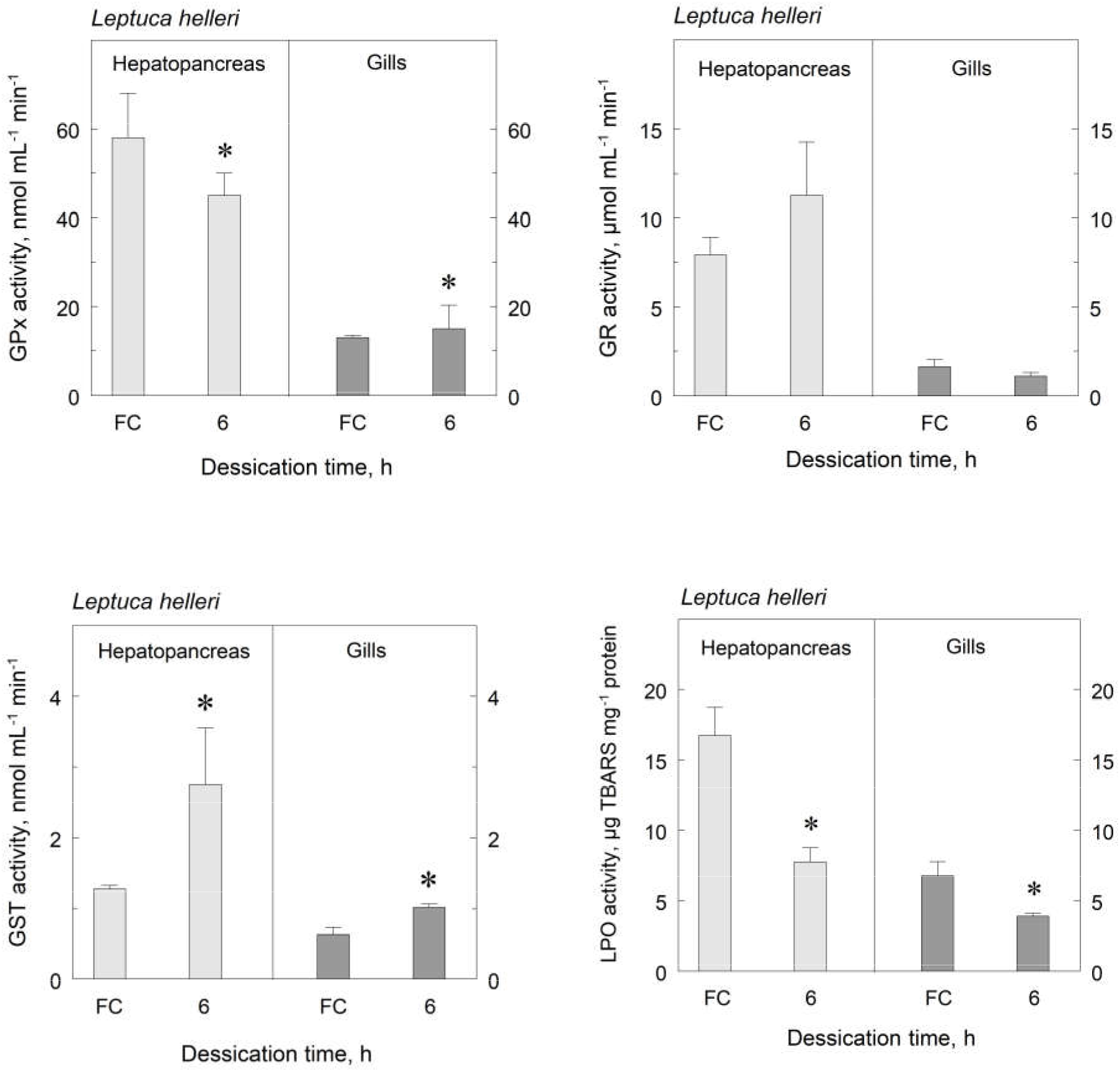
Enzyme activities (Glutathione peroxidase [GPx], Glutathione S-transferase [GST], Glutathione Reductase [GR] and Lipid peroxidation [LPO]) in hepatopancreas and gill homogenates from the endemic Galapagos fiddler crab *Leptuca helleri* kept emerged without access to water for 6 h after 2 days held at 29 ‰S (fresh caught crabs, FC). Data are the mean ± SEM (N=10). *P<0.05 compared to fresh caught crabs.

### Integrated biomarker response indexes

The response scores for each biomarker and the integrated biomarker response indexes are given in Figure 9 and associated tables.

**Figure 9.**
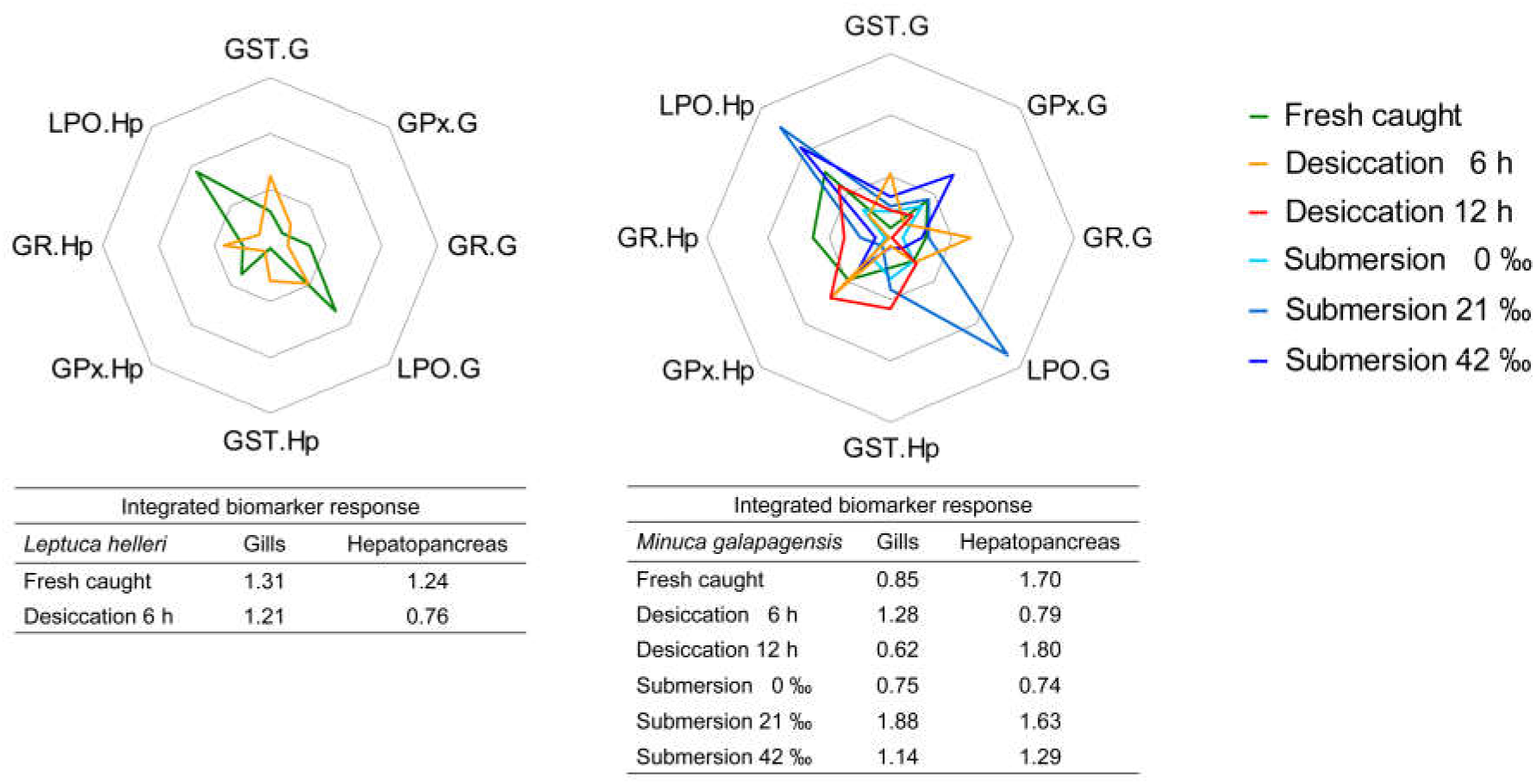
Integrated biomarker response indexes for the Galapagos fiddler crabs *Leptuca helleri* (left panel and table) and *Minuca galapagensis* (right panel and table) when maintained fully submerged for 6 h at salinities of 0 ‰S [distilled H_2_O, hypo-osmotic challenge], 21 ‰S [630 mOsm/kg H_2_O, isosmotic reference medium] or 42 ‰S [1,260 mOsm/kg H_2_O, hyper-osmotic challenge) after 2 days held at 29 ‰S (670 mOsm/kg H_2_O, fresh caught crabs), or held emerged without access to water for up to 12 h.

For *L. helleri* (Figure 9, left panel), fresh caught crabs showed the highest LPO indexes in both the gills and hepatopancreas (table below left panel). Desiccation for 6 h had little effect.

For *M. galapagensis*, submersion at 21 ‰S and desiccation for 6 h were the most relevant effectors of response (Figure 9, right panel). In the gills, LPO activity in 21‰S, and GST and GR activity after 6 h desiccation were the main determinants. In the hepatopancreas, 12 h desiccation and fresh caught crabs predominated as effectors (Figure 9 and table below right panel).

## Discussion

Our findings reveal clear differences in habitat characteristics, burrow densities, survival ability on submersion and desiccation challenge, osmoregulatory ability and oxidative stress enzyme behavior between the chosen populations of the two species of Galapagos fiddler crabs investigated on Santa Cruz Island.

The population of *Leptuca helleri* studied inhabits the intertidal zone where it encounters little variation in salinity and substrate moisture across the shore (Figure 3). Burrow densities, highest on the low shore, decline rapidly and markedly by 80% within just 10 meters, and enigmatically, cannot be attributed to variation in measured transect parameters. This population of *L. helleri* is abundant a fair distance from a mangrove stand where daily tidal submersion likely confers protection against water loss. In contrast, *M. galapagensis* inhabits the supralittoral zone, subject to only occasional seasonal tidal inundation, and is distributed across marked salinity and substrate moisture gradients. Burrow densities decline only slowly and progressively with increasing salinity and decreasing moisture (Figure 3), suggesting better salinity and desiccation tolerance than *L. helleri*. The burrows of *M. galapagensis* tend to be associated with mangrove vegetation, which together with thermoregulatory behavior (Smith and Miller, 1973) may alleviate temperature stress and evaporative water loss (McGuinness, 1994). This population of *M. galapagensis* is encountered farther from the nearest water source than *L. helleri* and may be more subject to water loss.

The two species exhibit striking differences in desiccation and submersion tolerances. *Leptuca helleri* cannot survive more than 6 h without water under experimental conditions, neither does it tolerate rigorous hypo- or hyper-osmotic challenge under forced submersion, surviving only at a salinity (21 ‰S) moderately dilute compared to burrow salinity (34 to 37 ‰S) and at which it hyper-regulates strongly. Forced submersion together with severe salinity challenge leads to death likely through a synergistic effect on osmoregulatory ability such as hypoxia, and insufficient energy available owing to a putative shift to anaerobic metabolism (Teal and Carey, 1967).

In contrast, *M. galapagensis* tolerated substantial experimental desiccation and submersion, showing no mortality. The crab hyper/hypo-regulates well, showing little if any effect of forced salinity submersion. *Uca rapax*, exposed to salinities between 40 and 63 ‰S (1,200 to 1,890 mOsm/kg H_2_O) hypo-regulates hemolymph osmolality between 1,069 and 1,085 mOsm/kg H_2_O; however, the submerged crabs osmoconform (Zanders and Rojas, 1996b) as also seen in *Uca pugilator* (D’orazio and Holliday, 1985). Forcibly submerged *M. rapax* shows a diminished ability to hypo-regulate hemolymph osmolality and [Na^+^] and [Cl^-^] at 60 ‰S (Capparelli et al., 2017), which may be due partly to limited oxidative ATP production (Teal and Carey, 1967).

Osmoregulatory ability in *M. galapagensis* was unaffected during submersion in roughly isosmotic or hyper-osmotic media (21 and 42 ‰S), although *L. helleri* could not survive severe hypo- or hyperosmotic challenge (0 and 42 ‰S) for 6 h. Anaerobic lactate metabolism predominates on submersion in fiddler crabs (Teal and Carey, 1967) and gill Na^+^/K^+^-ATPase activity increases in fiddler crabs submerged at low salinities (D’orazio and Holliday, 1985; Capparelli et al., 2017). *Leptuca helleri* may be unable to generate energy sufficient to osmoregulate and tolerate the synergic stress of forced submersion in severely hypo- and hyper-osmotic media.

Reactive oxygen species (ROS) are produced as result of normal cellular metabolism and can be induced by various environmental factors. They are highly reactive and damage molecules such as DNA, carbohydrates, lipids and proteins, altering their functions (Birben et al., 2012). Water flow through the gills is maintained during submersion, increasing O2 availability: ROS concentrations can subsequently increase, augmenting oxidative stress. The ROS defense system (GPx, GST and GR) in *M. galapagensis* is altered during submersion, together with increased oxidative stress (increased LPO levels) at 21 and 42 ‰S, particularly in the hepatopancreas, and at 21 ‰ S in the gills, possibly increasing tissue permeability (Birben et al., 2012). Elevated salinities generate oxidative stress and affect antioxidant mechanisms in fiddler crabs (Zanders and Rojas, 1996; Capparelli et al., 2017). Thus, salt secretion may induce metabolic shifts that generate ROS. Coherently, the integrated biomarker indexes for *M. galapagensis* reveal an increase in antioxidant defenses and oxidative stress in the gills of crabs at 21 and 42 ‰S and a decrease at 0 ‰S.

When crustaceans are exposed to desiccation, gas exchange may decrease since the gill lamellae collapse, reducing the diffusional surface area available (Withers, 1992; Morris and Oliver, 1999a), oxidative challenge consequently diminishing. In the gills, desiccation decreases the overall biomarker response in *L. helleri* and increases in *M. galapagensis*. Coherently, the integrated biomarker indexes show that antioxidant defense activity and oxidative stress diminish on desiccation in the hepatopancreas of both species.However, on longer aerial exposure, as seen for *M. galapagensis* after 12 h, the decreased oxygen supply can gill antioxidant defense activity manifest in the markedly reduced integrated biomarker response index, and severely affect energy balance and reserves, and may shorten survival time in aquatic animals (Martínez-Álvarez *et al*., 2005; Abele *et al*., 2007; Paital, 2013). Hemolymph volume in the crab *Lithodes santolla* decreases with desiccation, reducing oxygen titers and leading to systemic effects (Urbina *et al*., 2013). *Leptuca helleri* does not survive desiccation for much more than 6 h, when curiously, its hemolymph osmolality decreases. In contrast, *M. galapagensis* survives 12 h or more, its hemolymph osmolality increasing transiently, revealing water loss, returning incongruously to initial values.

Glutathione plays a major role in different cellular compartments, promoting elimination of xenobiotics and acting within the antioxidant system. It reacts directly with ROS, providing protective functions such as reduction, conjugation and interaction with other non-enzymatic antioxidants like vitamins E and C (Forman *et al*., 2019). Glutathione peroxidase (GPx) is responsible for scavenging organic and inorganic peroxides, glutathione S-transferase (GST) catalyzes the biotransformation of xenobiotics, and glutathione reductase (GR) reduces glutathione disulfide to glutathione (GSH), the sulfhydryl form, an important cellular antioxidant (Morris et al., 2014). The decreased hepatopancreas GPx and GR activities seen in *M. galapagensis* during submersion may conserve energy then made available during aerial exposure. Antioxidant enzymatic activities also decrease after 8 h aerial exposure in *Neohelice granulata* (de Oliveira *et al*., 2005). The glutathione system seems to be important during the emersion/submersion transition since all activities are elevated in fresh caught crabs, and GST and GR activities become much reduced during desiccation. This enzymatic defense system in the hepatopancreas and gills of *M. galapagensis* may prevent oxidative damage during desiccation, as suggested by the low lipid hydroperoxide activities.

The gills and hepatopancreas of the two fiddler crabs exhibited distinct oxidative responses. In *M. galapagensis*, forced submersion induced pronounced alterations in all biomarkers in the hepatopancreas. Oxidative stress during desiccation remained fairly unchanged or diminished in both tissues, with a likely reduction in ROS titers ensuing. In *L. helleri*, the oxidative stress system seems to be effective up to 6 h desiccation, with a decrease in oxidative stress titers (LPO) in both tissues. Such tissue-specific differences may be widespread since each responds differently to environmental parameters such as salinity challenge in the mud crab *Scylla serrata* (Paital and Chainy, 2010). This is corroborated by the integrated biomarker index, where each tissue exhibited a different score, based on treatment.

In conclusion, our findings reveal marked differences in tolerance of forced submersion and desiccation in the two fiddler crabs that inhabit the Galapagos Islands, *Minuca galapagensis* being much more resistant than *Leptuca helleri*, owing to its physiological and biochemical adjustments. *Minuca galapagensis* is a generalist species, manifesting few physiological and oxidative stress effects while the more ecologically demanding species, *Leptuca helleri*, cannot survive such conditions for long.

## Acknowledgements

We wish to express our appreciation to the staff of the regional offices of the Ministerio del Ambiente de Ecuador (MAE) for supporting this research, and in particular, we thank Galo Quezada and Jeniffer Suarez (Galapagos National Park, Isla Santa Cruz, Permit #083-2019 DPNG). During fieldwork, we were graciously assisted by Angel Cajas (Ikiam), Alexandra Kler Lago (Galapagos National Park) and Mara Anais Espinoza Buitrón (Galapagos National Park). MVC received no financial support towards the costs of this investigation. The University of Northern Iowa (UNI) Study Abroad Program, and Information Technology Services, provided travel support for CLT. JCM received an Excellence in Research scholarship from the Conselho Nacional de Desenvolvimento Científico e Tecnológico, Brazil (CNPq 303613/2017-3) which defrayed travel and accommodation costs. PKG-C thanks the Fundação de Amparo à Pesquisa do Estado de São Paulo, Brazil for financial support (FAPESP #2017/04970-5). We are also grateful to Dr. Gabriel Massaine Moulatlet (Ikiam) for preparation of the study area map and Figure 3.

## Author contributions

*Mariana Vellosa Capparelli*: Conceptualization, Methodology, Validation, Formal analysis, Investigation, Resources, Writing - Original Draft, Writing - Review & Editing, Supervision, Project administration. *Carl Leo Thurman*: Methodology, Resources, Writing - Review & Editing, Funding acquisition. *Paloma Gusso Choueri*: Validation, Writing - Review & Editing. *Denis Moledo Abessa*: Validation, Resources, Writing - Review & Editing. *Mayana Karoline Fontes*: Validation, Writing - Review & Editing. *Caio Rodrigues Nobre*: Software, Formal analysis, Writing - Review & Editing. *John Campbell McNamara*: Methodology, Validation, Formal analysis, Investigation, Resources, Data curation, Writing - Original Draft, Writing - Review & Editing, Supervision, Project administration, Funding acquisition.

## Compliance with Ethical Standards

This study complies with all Ecuadorian, Brazilian, institutional and international guidelines on the use of invertebrate animals in scientific research.

